# EEG alpha-theta dynamics during mind wandering in the context of breath focus meditation: an experience sampling approach with novice meditation practitioners

**DOI:** 10.1101/2020.10.23.351759

**Authors:** Julio Rodriguez-Larios, Kaat Alaerts

**Author notes:** **Corresponding author: Julio Rodriguez-Larios**, Research Group for Neurorehabilitation (eNRGy), Tervuursevest 101 - box 1501, 3001 Leuven (Belgium).

## Abstract

Meditation practice entails moments of distraction dominated by self-generated thoughts (i.e. mind wandering). Initial studies assessing the neural correlates of mind wandering in the context of meditation practice have identified an important role of theta (4-8 Hz) and alpha (8-14 Hz) neural oscillations. In this study, we use a probe-caught experience sampling paradigm to assess spectral changes in the theta-alpha frequency range during mind wandering in the context of breath focus meditation. Electroencephalography (EEG) was measured in 25 novice meditation practitioners during a breath focus task in which they were repeatedly probed to report whether they were focusing on their breath or thinking about something else. Mind wandering episodes were associated with an increase in the amplitude and a decrease in the frequency of theta (4-8 Hz) oscillations. Conversely, alpha oscillations (8-14 Hz) were shown to decrease in amplitude and increase in frequency during mind wandering relative to breath focus. In addition, mind wandering episodes were shown to be accompanied by increased harmonicity and phase synchrony between alpha and theta rhythms. Because similar spectral changes in the theta-alpha frequency range have been reported during controlled cognitive processes involving memory and executive control, we speculate that mind wandering and controlled processes could share some neurocognitive mechanisms. From a translational perspective, this study indicates that oscillatory activity in the theta-alpha frequency range could form adequate parameters for developing EEG-neurofeedback protocols aimed at facilitating the detection of mind wandering during meditation practice.

## INTRODUCTION

In the last years, secularized versions of Buddhist contemplative practices have become increasingly common in our society due to their putative health benefits. An important part of these practices consist of directing the attentional focus to a specific object (e.g. the sensation of breathing) while ‘noticing’ and ‘letting pass’ self-generated thoughts (i.e. mind-wandering episodes) (Anālayo, 2019; Delorme & Brandmeyer, 2019; Hanh, 1990; Krishnamurti, 2002). In this type of meditation practice (i.e. ‘focused attention meditation’; see Lutz et al., 2015) periods of focused attention to the object of meditation are alternated with the occurrence of mind wandering episodes involving memories, future plans, and fantasies (Schooler et al., 2014; Smallwood & Schooler, 2015).

Only a few studies have directly assessed the EEG correlates of mind wandering during meditation practice. For this purpose, two main paradigms have been adopted: self-caught experience sampling (i.e. participants press a button whenever mind wandering is noticed) and probe-caught experience sampling (i.e. probing participants at random time intervals throughout the meditation to report whether they were mind wandering). Three previous studies adopting experience sampling paradigms have shown that mind wandering episodes during meditation are associated with decreased alpha power while mixed results were reported in the theta range (Braboszcz & Delorme, 2011; Brandmeyer & Delorme, 2018; van Son et al., 2019). These results are in line with previous EEG studies that compared meditation practice with resting state and inferred that the reported changes (increased alpha amplitude during meditation is the most consistent finding; see Cahn & Polich, 2006; Lee et al., 2018; Lomas et al., 2015) could be partially explained by reduced mind wandering during meditation (Faber et al., 2015; Hinterberger et al., 2014; Lehmann et al., 2012).

In recent years, the EEG research field has witnessed an increased emphasis on studying interactions among neural rhythms (i.e. cross-frequency coupling), rather than studying distinct rhythms in isolation (Canolty & Knight, 2010). In this way, communication among brain rhythms is thought to be implemented via cross-frequency phase synchrony, as it is the only form of cross-frequency coupling that allows consistent neuronal spike-time relationships (Fell & Axmacher, 2011; Palva & Palva, 2017). In this context, a recent theory posited that interactions among brain rhythms at different frequencies are enabled by the formation of harmonic cross-frequency arrangements (Klimesch, 2013, 2018). The rationale behind Klimesch (2013, 2018) theory is that phase synchrony between two rhythms that oscillate at different frequencies is only mathematically possible when they form a harmonic cross-frequency ratio (e.g. 10 Hz and 5 Hz in the case of a 2:1 relationship between alpha and theta rhythms). Therefore, when interactions between two brain rhythms are required, their center frequency would need to shift in order to form a harmonic cross-frequency ratio and this way enable their information exchange via cross-frequency phase synchronization.

In the light of the theoretical framework proposed by Klimesch (2013, 2018), two recent studies suggested an important role of alpha-theta cross-frequency dynamics in meditation practice (Rodriguez-Larios, Faber, et al., 2020; Rodriguez-Larios, Wong, et al., 2020). Specifically, in a recent study with highly experienced meditatiors, it was shown that meditaitve states aimed at achieving a state of ‘mental emptiness’ was associated with a reduction in alpha:theta phase synchrony / harmonicity when compared to non-meditative resting state (Rodriguez-Larios, Faber, et al., 2020). In the same line, it was shown that an 8-week mindfulness training in novices (hours of practice and attendance to a mindfulness course) was associated with reduced alpha:theta phase synchrony / harmonicity during focused attention meditation (Rodriguez-Larios, Wong, et al., 2020). Since meditation training has been associated with reduced mind wandering during meditation practice (Brandmeyer & Delorme, 2018), it was postulated that the observed reductions in transient alpha-theta phase synchrony / harmonicity are reflective of a reduction in mind wandering episodes (Rodriguez-Larios, Faber, et al., 2020; Rodriguez-Larios, Wong, et al., 2020). In this way, it can be anticipated that, due to the putative role of alpha and theta rhythms in memory and executive control (Akiyama et al., 2017; de Vries et al., 2020; Rodriguez-Larios & Alaerts, 2019; Schack et al., 2005), a certain degree of alpha-theta phase synchrony is necessary for the generation of mind wandering episodes, as these would require the integration between memory and executive components of cognition (Kam & Handy, 2014; Smallwood & Schooler, 2006).

In this study, we used a probe caught experience sampling paradigm to characterize EEG alpha-theta dynamics during mind wandering in the context of meditation practice. To do so, 25 novice meditation practitioners were asked to perform a breath focus meditation task in which they were frequently probed to report whether they were focusing on their breath or thinking about something else. Based on previous results (Rodriguez-Larios, Faber, et al., 2020; Rodriguez-Larios, Wong, et al., 2020), we hypothesized that self-reported episodes of mind wandering (relative to breath focus) would be associated with spectral changes in the theta-alpha frequency range that would facilitate harmonicity and cross-frequency phase synchrony between alpha (8 – 14Hz) and theta (4-8 Hz) rhythms.

## METHODS

### Participants

Twenty-eight subjects (11 males, mean age 23.46 years, age range: 20-29) participated in the study. Participants were recruited through flyers and social media from various campuses of the University of Leuven. Informed written consent was obtained from all participants before the study. Consent forms and study design were approved by the Social and Societal Ethics Committee (SMEC) of the University of Leuven, in accordance to the World Medical Association Declaration of Helsinki (dossier no. G- 2018 12 1463). Participants were compensated for their participation (8 € per hour). Three participants were excluded due to technical problems during data acquisition.

### Design and task

EEG recordings were obtained while participants performed a breath focus task (Figure 1). In this task, participants were instructed to focus on the sensation of breathing while closing their eyes. At random intervals (20 to 60 seconds), the participants were presented with a bell sound and were required to open their eyes and respond to the following questions: *(i) “What were you doing right before you heard the sound?”* (two response options: focusing on the sensation of breathing or thinking about something else) *(ii) “How confident are you of your answer?”* (7-point Likert scale; 1: not confident at all; 7: completely confident)*? “How were you feeling just before you heard the sound?”* (5-point Likert scale based on the self-assessment Manikin rating of arousal; 1: excited; 5: calm; Bradley & Lang, 1994) (see Figure 1). The instructions of the task and the questions were displayed on the computer screen in front of the participants (programmed with E prime 2.0 software). Prior to the experimental procedure, participants performed one practice trial. The task consisted of a total of 40 trials (total duration of approximately 40 minutes). After completion of the task, debriefings were performed in which participants were asked to report their level of drowsiness with the question *“How often did you feel like falling asleep during the task?”* on a 7-point Likert scale (1=never, 2=very rarely (~15%), 3=rarely (~30%), 4 = occasionally (~50%), 5 = often (~70%), 6 = very often (~85%) and 7= all the time (100%)). Also debriefings of the participants’ level of arousal and emotional valence were assessed retrospectively after the task using 5-point Likert scales based on the self-assessment Manikin ratings (Bradley & Lang, 1994) (1: arousing/unpleasant; 5: calm/pleasant). Note that these debriefings of arousal, valence and drowsiness were only available for 22 (out of 25) participants. Furthermore, the breath focus task was part of a larger assessment protocol consisting of several other experimental conditions that are not part of the current report: i.e., a rest condition (preceding the breath focus task) and an unguided meditation session, a heartbeat counting task and an arithmetic task (performed after the breath focus task).

**Figure 1.**
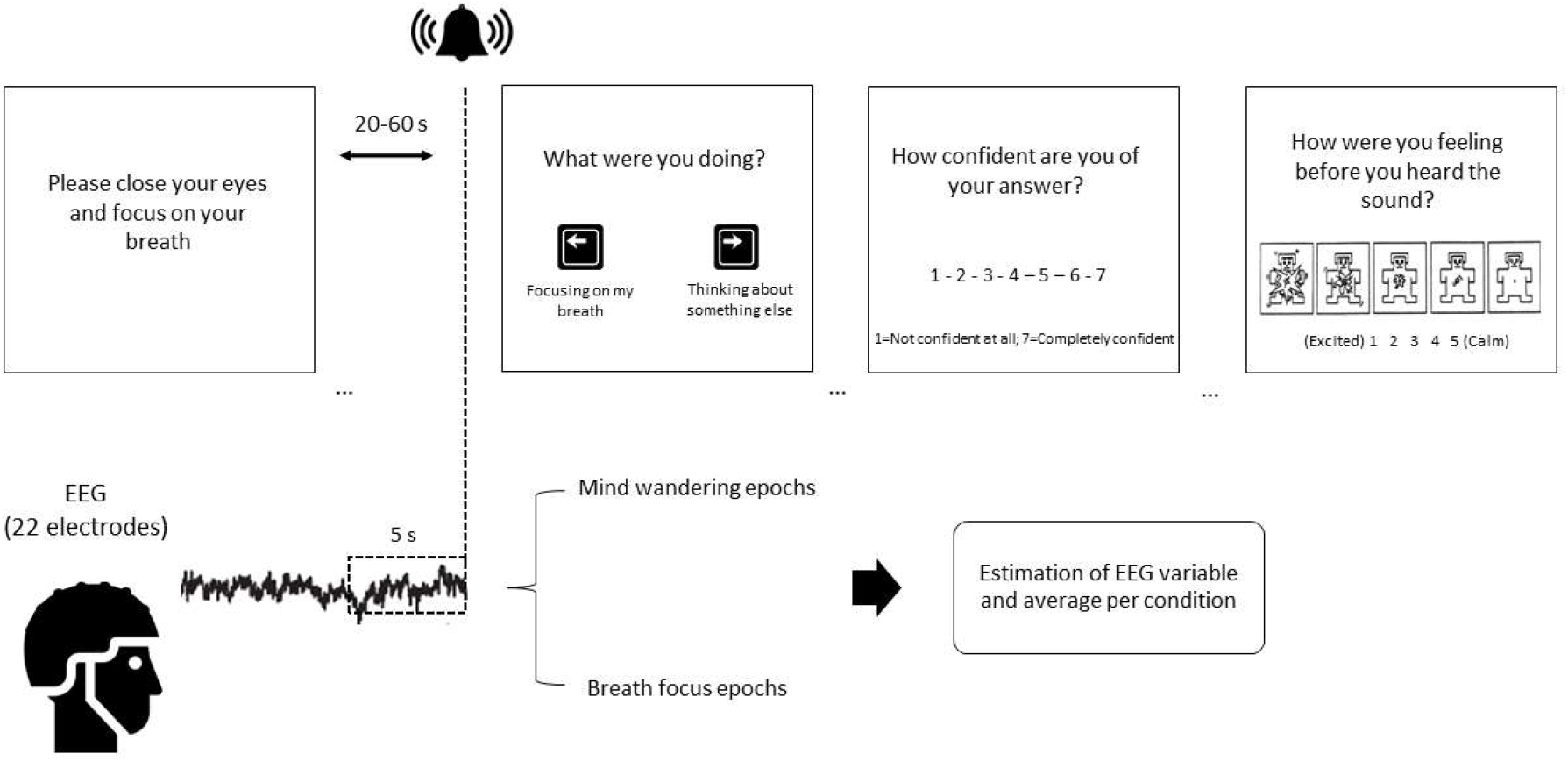
Depiction of experimental design and analytical approach. Participants were instructed to focus on their breath while keeping their eyes closed and to open their eyes every time they heard a bell sound. After the bell sound, participants had to report whether they were focusing on their breath or thinking about something else using a computer’s keyboard. Additional questions involving self-ratings of their level of confidence and arousal were also included in each trial. Participants completed 40 trials and the length of each trial varied between 20 and 60 seconds. Only the last 5 seconds before the bell sound were used for subsequent EEG analysis. These EEG epochs were sorted into the conditions breath focus (BF) or mind wandering (MW) depending on participant’s subsequent answer. Each EEG dependent variable was estimated per epoch an averaged across epochs from the same condition.

### EEG data acquisition and analysis

#### Recordings

The Nexus-32 system (version 2015a, Mind Media, The Netherlands) and BioTrace software (Mind Media) were used to perform electroencephalography (EEG) recordings. Continuous EEG was recorded with a 22 electrode cap (two reference electrodes and one ground electrode) positioned according to the 10-20 system (MediFactory). Vertical (VEOG) and horizontal (HEOG) eye movements were recorded by placing pre-gelled foam electrodes (Kendall, Germany) above and below the left eye (VEOG) and next to the left and right eye (HEOG) (sampling rate of 2048 Hz). Skin abrasion and electrode paste (Nuprep) were used to reduce the electrode impedances during the recordings. The EEG signal was amplified using a unipolar amplifier with a sampling rate of 512 Hz. EEG recordings were synchronized to the presented task using E-prime and the Nexus trigger interface (Mind Media).

#### Pre-processing

Pre-processing was performed using custom MATLAB scripts and EEGLAB functions (Delorme & Makeig, 2004). EEG data was first filtered between 1 and 40 Hz (function *pop_eegfiltnew*). Abrupt noise in the data was removed using the Artifact Subspace Reconstruction method (function *clean_asr* with a cut-off value of 20 SD; see Chang, Hsu, Pion-Tonachini, & Jung, 2020). Then, data were re-referenced to average electrodes and epoched to −5 seconds relative to the bell sound probes. Then, epochs with absolute amplitudes exceeding 100 μV were excluded from further analyses (i.e. 7 epochs across all participants; 5 mind wandering epochs and 2 breath focus epochs). Independent Component Analysis (ICA) on concatenated data (across conditions) was performed to correct for eye movements. Components were rejected based on their spatial topography and their correlation with H/VEOG electrodes. ICA-based eye movement correction led to an average removal of 1.68 ± SD 0.47 components per subject. Finally, clean epochs of EEG data were sorted in the categories breath focus (BF) or mind wandering (MW) depending on subject’s subsequent answers.

#### Alpha-theta amplitude estimation

Short term fast Fourier transformations were performed using the function *spectrogram* in MATLAB 2019b (sliding window = 1 second; overlap = 90%; 0.5 Hz resolution between 4 and 14 Hz). Both absolute (μV) and relative (% of total power) amplitudes were averaged across the 5 seconds epochs per subject and electrode.

#### Cross-frequency ratios, mean frequency and peak detectability estimation with the find peaks (FP) approach

Short time Fourier transformations were applied to the pre-processed EEG data using the *spectrogram* function implemented MATLAB r2019b (sliding window = 1 second; overlap = 90%) to compute the time varying spectrum between 4 and 14 Hz. The signal in each sliding window was zero padded to obtain smoother frequency spectra and this way improve peak detection (i.e. 0.1 Hz resolution) (Haegens et al., 2014). Transient peak frequencies in the alpha (8-14Hz) and theta (4-8Hz) frequency bands were defined using the MATLAB function *findpeaks*, which define peaks as data samples that are larger than its two neighboring samples. When more than one peak was detected the peak with the highest amplitude was selected. The find peaks algorithm detected at least one peak in 99.90% (SD = 0.05) of the time points in the alpha band (8-14 Hz) and in 95.11% (SD = 4.96) of the time points in the theta band (4-8 Hz). Note that also additional analyses were performed using stricter criteria for defining alpha and theta peaks (i.e. amplitude threshold based on the 1/f trend of the frequency spectrum). The 1/f trend of the spectrum was estimated per subject and electrode (averaged across trials) by fitting a straight line in log–log space to the EEG frequency spectrum (1 to 30 Hz excluding the alpha peak) using the *robustfit* function in MATLAB (for similar approaches see Caplan et al., 2015; Kosciessa et al., 2020; Watrous et al., 2018; Whitten et al., 2011). When using an amplitude threshold based on the 1/f trend of the spectrum, the find peaks algorithm detected at least one peak in 95.20% (SD = 9.29) of the time points in the alpha band and in 76.28 % (SD = 15.48) of the time points in the theta band.

The identified transient peak frequencies in the alpha and theta bands (with or without amplitude threshold based on the 1/f trend) were used to compute their numerical ratio per time point. These transient cross-frequency ratios were averaged to the first decimal place in order to estimate the proportion (%) of time points showing each possible cross-frequency ratio (from 1.1 to 3.4). Proportions of cross-frequency ratios, computed for each epoch, were subsequently averaged across epochs from the same condition (BF or MW) per subject and electrode. In particular, a weighted average was calculated based on the confidence level of each answer (i.e. ranging from 1 to 7), such that trials with a higher level of confidence were weighted higher in the total average score.

In addition, to explore condition-related differences in the mean frequencies of the alpha and theta bands, transiently detected peak frequencies were averaged across time points (within an epoch) and across epochs from the same condition (also weighted for confidence) separately for each subject, electrode and frequency band. Finally, to assess possible condition-related differences in peak detectability, the percentage of epochs in which *no* alpha or theta peaks were detected (with and without amplitude threshold based on the 1/trend of the spectrum) was computed for each epoch and averaged (weighted for confidence) across epochs from each condition (BF or MW) separately for each subject and electrode.

#### Cross-frequency ratios and mean frequency estimation with the instantaneous frequency (IF) approach

In addition to the find peaks approach, we used an alternative method for defining transient cross-frequency ratios and the mean frequencies of alpha and theta rhythms. Namely, the instantaneous frequencies within the alpha (8–14 Hz) and theta (4–8 Hz) bands were computed using the method and MATLAB code developed by Cohen (2014). This method estimates transient changes in the frequency of an oscillator based on the temporal derivative of the phase angle time series. For this purpose, the EEG signal was first filtered using a plateau-shaped zero phase digital filter (MATLAB function *firl1*). Next, the Hilbert transform was applied to extract the phase angle time series based on the analytical signal. The instantaneous frequency was calculated by multiplying the first temporal derivative of the phase angle by the sampling rate and dividing it by 2π. Finally, a median filter was applied to the instantaneous frequency time series (10 steps between 10 and 400 ms) in order to attenuate non-physiological frequency jumps.

Similar to the find peaks approach, the numerical ratio between transient alpha and theta (instantaneous) frequencies was computed per time point (within an epoch) in order to estimate the proportion (%) of time points showing each possible cross-frequency ratio (from 1 to 3.5) per epoch. Also here, in order to assess condition-related differences, proportions of cross-frequency ratios were averaged (weighted by confidence ratings) across epochs from the same condition (MW or BF) separately for each subject and electrode. Further, to avoid the possibility of estimating instantaneous frequencies (and therefore cross-frequency ratios) in the absence of actual peaks (Aru et al., 2015), we performed an additional analysis in which only trials with observable alpha and theta peaks above the estimated 1/f trend of the spectrum (see previous section) (i.e., averaged for each 5-second epoch) were selected. Accordingly, in this additional analysis, the calculation of alpha-theta cross-frequency ratios was performed for an average of 13.72 ± SD 6.45 (BF) and 11.27 ± SD 6.13 (MW) trials (estimation across electrodes and subjects).

In addition, similar to the find peaks approach, condition-related differences were also explored for alpha and theta mean frequency. For this purpose, the identified instantaneous frequencies were averaged within epochs and across epochs from the same condition (MW or BF) separately for each subject and electrode. This analysis was also performed with the previously described selection of trials (i.e. only trials with observable alpha and theta peaks above the estimated 1/f trend of the spectrum).

#### Cross-frequency phase synchrony

Phase synchrony was estimated using the n:m phase locking value (PLV) (sliding window = 300ms) which quantifies the consistency of phase relationships between two oscillators (Palva et al., 2005; Palva & Palva, 2017; Siebenhühner et al., 2016). For that purpose, the instantaneous phase of alpha and theta rhythms were obtained after the application of a FIIR filter and Hilbert transform. Then, the instantaneous phase difference between alpha and theta rhythms was calculated to create unitary vectors whose mean length is proportional to the level of phase synchronization (Tass et al., 1998). Within the given frequency bands definition, phase synchrony between alpha (8 - 14Hz) and theta (4 - 8Hz) rhythms is only possible for the cross-frequency ratios 2:1 and 3:1. Accordingly, the n:m phase-locking value was estimated with n=1 and m=2 or 3. PLV was estimated in each 5-seconds epoch per subject and electrode. In order to avoid a spurious estimation of phase (Aru et al., 2015), we performed an additional analysis in which PLV was only computed in those trials in which alpha and theta peaks were detected above the estimation of the 1/f trend of the spectrum (the 1/f trend of the spectrum was estimated as previously described).

### Data handling and statistical analysis

#### Behaviour and self-reports

Trial-specific debriefings of arousal and confidence obtained during the task were averaged across trials of the BF and MW condition (per subject) and paired-samples t-tests were used to assess condition-related differences. Additionally, Pearson correlation analyses were performed to assess the relationship between inter-individual variability in the proportion of mind wandering epochs (%) and general debriefings of participants’ level of drowsiness, arousal and emotional valence.

#### EEG

For each of the computed EEG dependent variables a cluster-based permutation statistical method was adopted to assess condition-related differences (between MW and BF) as implemented in the MATLAB toolbox *Fieldtrip* (function *ft_timelockstatistics*) (Maris & Oostenveld, 2007). This statistical method controls for the type I error rate arising from multiple comparisons through a non-parametric Montecarlo randomization. In short, data were shuffled (1000 permutations) to estimate a “null” distribution of effect sizes based on cluster-level statistics (sum of t-values with the same sign across adjacent electrodes and/or frequencies/ratios). Then, the cluster-corrected p value was defined as the proportion of random partitions in the null distribution for which the test statistics exceeded the one obtained for each significant cluster (cluster-defining threshold: p < 0.05) in the original (non-shuffled) data. Significance level for the cluster permutation test was set to 0.025 (corresponding to a false alarm rate of 0.05 in a two-sided test) (Maris & Oostenveld, 2007). To assess condition-related differences, paired-samples t-test (function *ft_statfun_depsamplesT* in Fieldtrip) was chosen as the test statistic. Additionally, repeated measures one-way ANCOVAs (as implemented IBM SPSS Statistics 27) were performed on the identified significant clusters (averaged within positive and within negative clusters separately) of each EEG dependent variable to assess whether the condition effect (MW-BF) was significantly influenced by drowsiness levels (reported through debriefings at the end of the task). Data was plotted using custom MATLAB scripts in combination with EEGLAB (Delorme & Makeig, 2004) and ‘raincloud’ (Allen et al., 2019) functions.

## RESULTS

### Self-reported mind wandering

Participants reported to be mind wandering in 43.5% (SD = 19.57) of the trials during the breath focus meditation task. No significant difference was revealed between the percentage of MW and BF trials (t (24) = 1.66; p = 0.10). Considering that the bell sound occurred at a variable time interval ranging between 20 to 60 seconds, we formally explored whether the time interval preceding the probe differed significantly between trials categorized as MW versus BF. However, no significant difference was found (t (24) = −1.15, p = 0.25), thereby indicating that the length of the time interval did not affect the probability of reporting MW or BF (e.g., trials with a longer time interval were not categorized more frequently as MW or BF trials).

### Condition-related amplitude modulations in the theta-alpha frequency range (4-14Hz)

Permutation statistics revealed a relative amplitude *increase* in the theta range (frequencies between 4 and 7.5 Hz; t _cluster_ (24) = 551.29; p <0.001) and a relative amplitude *decrease* in the alpha range (frequencies between 8 and 12.5 Hz; t _cluster_ (24) = −399.25; p = 0.002) during MW compared to BF (see Figure 2A). A qualitatively similar pattern of results was obtained when analyses were performed using absolute amplitude, although in this case, the condition effect in the theta range did not reach statistical significance (theta range 4-7 Hz: t _cluster_ (24) = 98.76; p = 0.058; alpha range 8-13 Hz: t _cluster_ (24) = −454.85; p < 0.001) (see Figure 2B).

**Figure 2.**
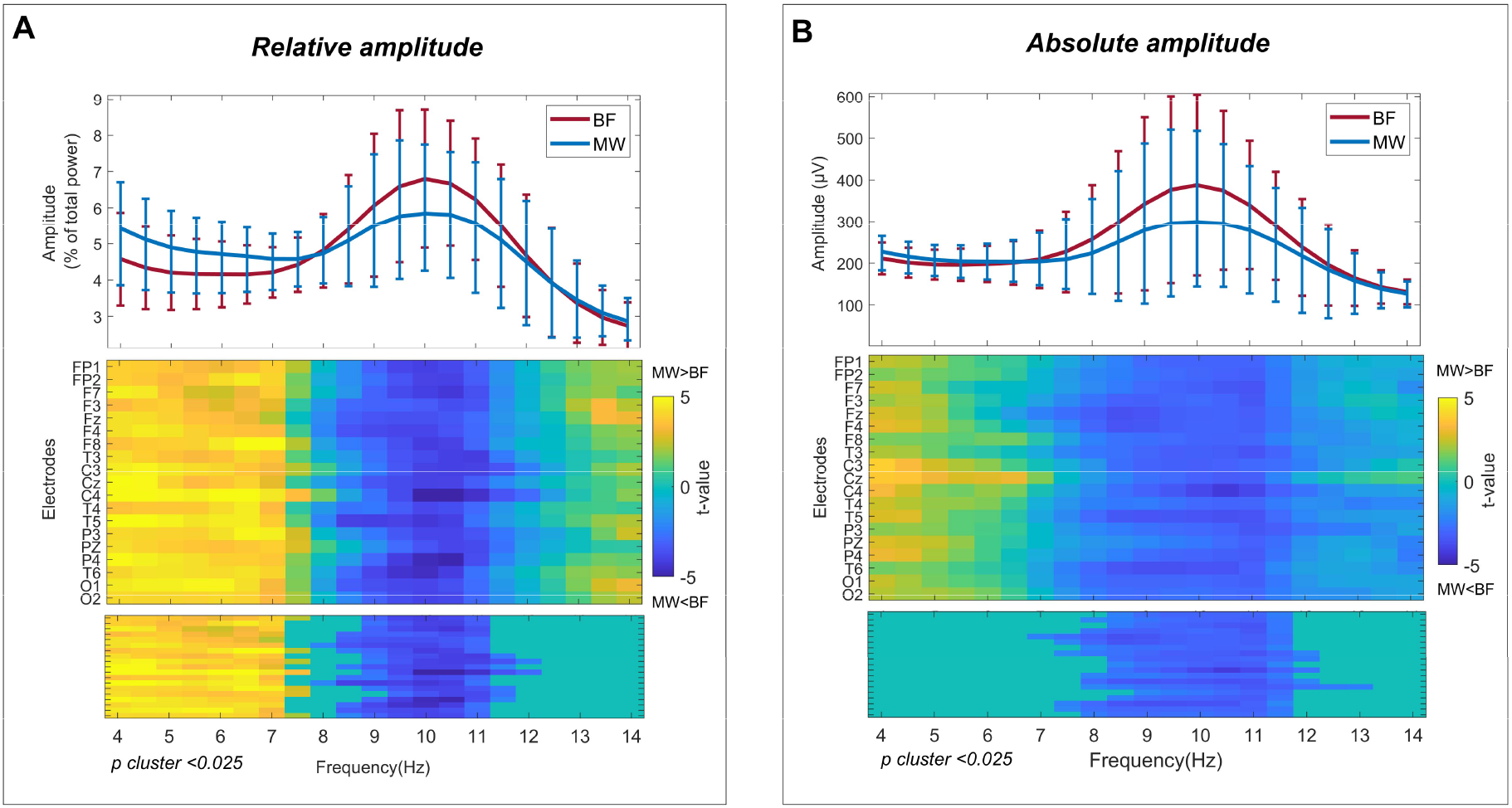
Condition-related amplitude modulations in the theta-alpha frequency range (4-14 Hz). Depiction of grand averages and condition effects (mind wandering (MW) – breath focus (BF)) with relative amplitude (Panel A) and absolute amplitude (Panel B) spectral estimations. The top panels show the mean amplitude per condition (MW in blue and BF in red) and frequency bin (4 to 14 Hz in 0.5 Hz bins) across electrodes and subjects (error bars represent standard deviation). Middle and bottom panels show t-values (resulting from paired samples t-tests) per frequency bin and electrode before and after correcting for multiple comparisons (cluster permutation correction in bottom plot; p-value < 0.025). MW (relative to BF) was associated to amplitude increases in the ~4-8 Hz frequency range (trend-level for absolute amplitude) and amplitude decreases in the ~8-14 Hz frequency range.

### Condition-related modulations in alpha and theta peak detectability

The probability of detecting at least one alpha peak was not significantly different between the MW and BF conditions (no significant clusters identified, see Supplementary Figure 1, panel A). A significant condition-related difference was evident however for the detection of theta peaks, indicating a *higher* probability of detecting a theta peak during the MW compared to the BF condition (t _cluster_ (24) = −9.53; p = 0.016) (Supplementary Figure 1, panel B). A qualitatively similar pattern of results was observed for theta peak detectability when using an amplitude threshold based on the 1/f trend (t cluster (24) = −5.73; p = 0.012) (see Supplementary Figure 1, panel D). However, note that in this case, also a condition effect for the detection of alpha peaks emerged, indicating a *lower* probability of detecting an alpha peak above the 1/f trend in the MW condition compared to the BF condition (t _cluster_ (24) = 53.27; p < 0.001) (see Supplementary Figure 1, panel C).

### Condition-related modulations in mean theta (4-8 Hz) and alpha (8-14 Hz) frequency

#### Instantaneous frequency approach

Permutation statistics revealed a higher mean alpha frequency (t _cluster_ (24) = 8.11, p = 0.024) and a lower mean theta frequency (t _cluster_ (24) =−41.98; p <0.001) during MW relative to BF (see Figure 3A-B). A qualitatively similar pattern of results was obtained when repeating the analysis with a selection of trials (only those showing alpha and theta peaks above the 1/f trend) (alpha: t _cluster_ (24) = 5.54, p = 0.027) (theta: t _cluster_ (24) = −4.92, p = 0.036) (see Supplementary Figure 2A-B).

**Figure 3.**
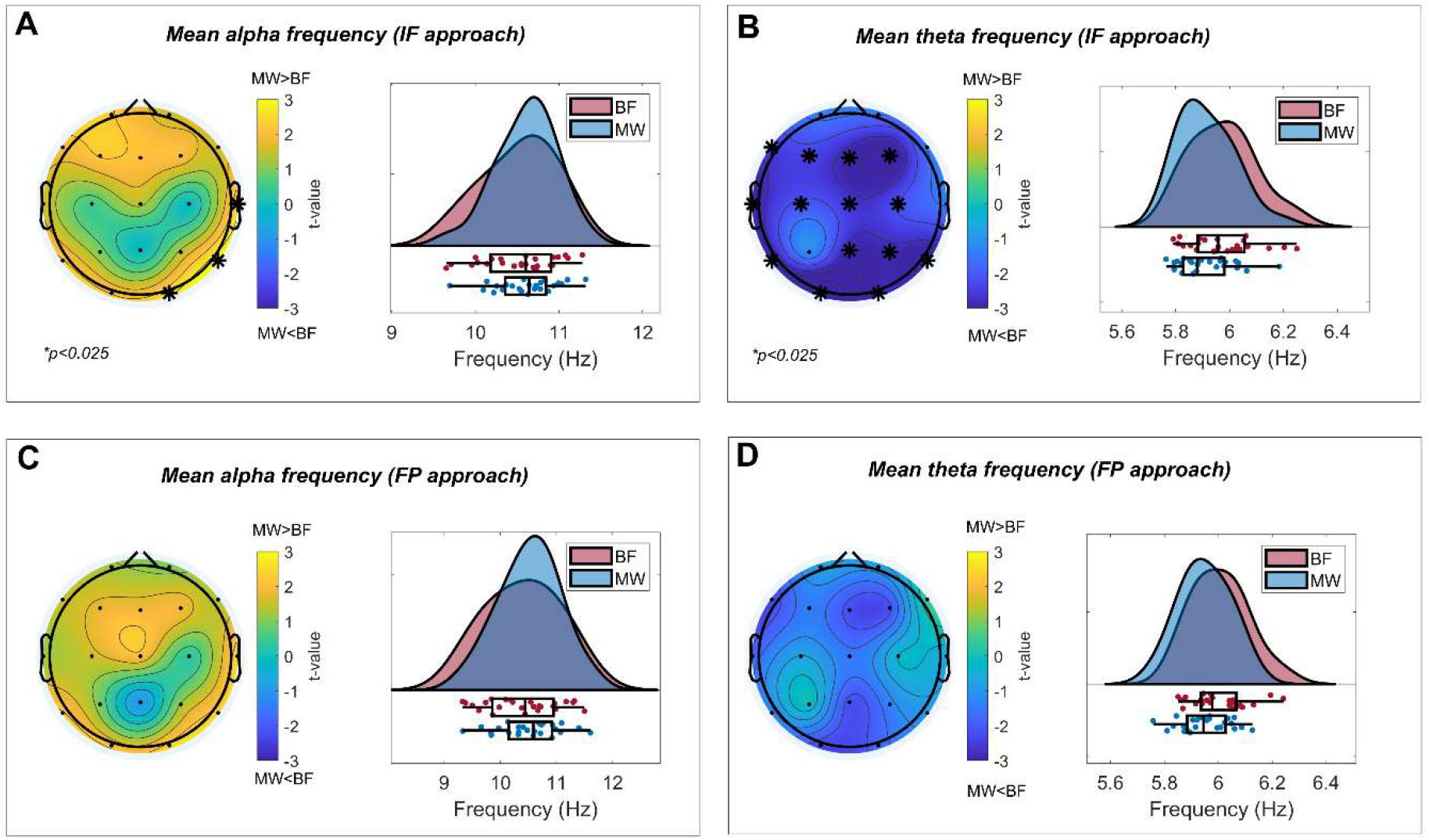
Condition-related modulations of alpha and theta mean frequency. Each panel depicts topographical maps representing condition-related changes (mind wandering (MW) - breath focus (BF)) in mean alpha and theta frequency per each electrode (left plot; t-values of paired samples t-test are color-coded) and the distributions of the identified clusters per condition (right plot; individual points are subjects). Mean frequency in alpha (8-14 Hz) and theta (4-8 Hz) bands were estimated through instantaneous frequency (IF) and find peaks (FP) analytical approaches. When estimated with the instantaneous frequency (IF) approach, MW was associated to higher mean alpha frequency (Panel A) and lower mean theta frequency (Panel B) compared to BF. A qualitatively similar pattern of results could be observed when mean frequencies were estimated with the FP approach (Panel C and D), although in this case, these changes only reached statistical tendency.

#### Find peaks approach

Permutation statistics revealed a statistical trend indicating higher mean alpha frequency (t _cluster_ (24) = 7.25, p = 0.034) and a lower mean theta frequency (t _cluster_ (24) = −4.47, p = 0.069) during MW relative to BF (see Figure 2C-D). A similar pattern of results was obtained when using the 1/f trend as an amplitude threshold for the detection of transient peaks, although note that here, condition-related differences were more pronounced (alpha: t _cluster_ (24) = 9.26, p = 0.019) (theta: t _cluster_ (24) = −15.28, p = 0.003) (see Supplementary Figure 2C-D).

### Condition-related modulations in the transient occurrence of alpha:theta cross-frequency ratios

#### Instantaneous frequency approach

Permutation statistics revealed an increased occurrence of alpha:theta ratios between 1.9 and 2.8 (t _cluster_ (24) = 153.67; p = 0.002) and a decreased occurrence of alpha:theta ratios between 1.2 and 1.6 (t _cluster_ (24) = −48; p = 0.021) during MW relative to BF when estimated with the IF approach (see Figure 4A). A qualitatively similar pattern of results was obtained when repeating the analysis with a selection of trials (i.e. trials showing alpha and theta peaks above the 1/f trend of the spectrum) (t _cluster_ (24) = 71.58; p < 0.001; t _cluster_ (24) = −24.16; p = 0.022) (see Supplementary Figure 3A).

**Figure 4.**
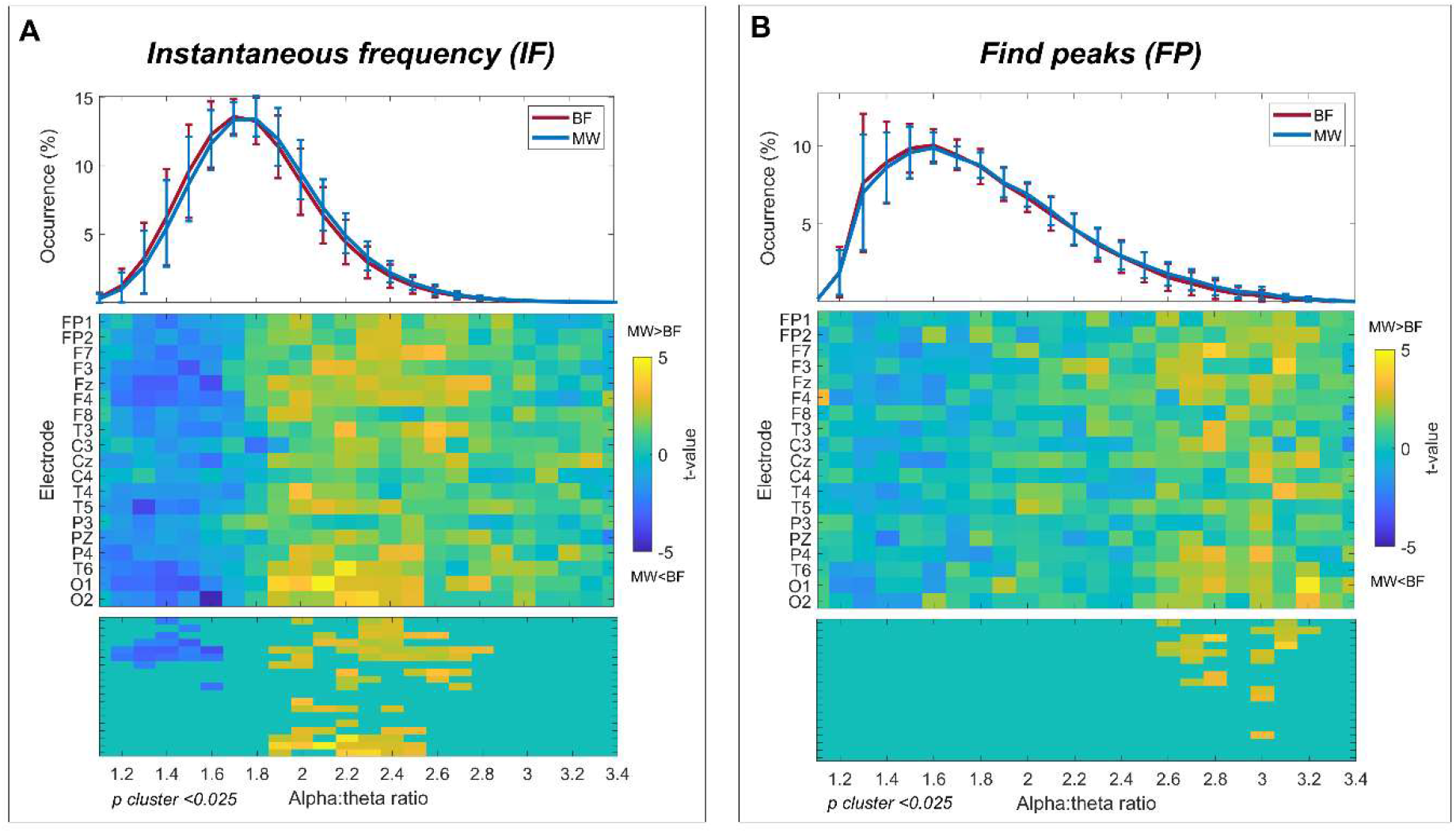
Condition-related modulations of transient alpha:theta cross-frequency ratios. Depiction of grand averages and condition effects (mind wandering (MW) – breath focus (BF)) in the occurrence of alpha:theta cross-frequency ratios when estimated with the instantaneous frequency (IF) (Panel A) and find peaks (FP) (Panel B) analytical approaches. The top panels show the mean occurrence of each alpha:theta ratio (from 1.1 to 3.4 in steps of 0.1 Hz) per condition (MW in blue and BF in red) across electrodes and subjects (error bars represent standard deviation). Middle and bottom panels show t-values (resulting from paired samples t-tests) per cross-frequency ratio and electrode before and after correcting for multiple comparisons (cluster permutation correction in bottom plot; p-value of cluster < 0.025). When estimated through IF, MW (relative to BF) was associated to an increase in the occurrence of alpha:theta ratios between 1.9 and 2.8 and a decrease in the occurrence of alpha:theta ratios between 1.2 and 1.6. For the FP approach, only a significant increase in the occurrence of alpha:theta ratios between 2.6 and 3.2 was reported.

#### Find peaks approach

Permutation statistics revealed an increased occurrence of alpha:theta ratios between 2 and 2.8 (t _cluster_ (24) = 12.17, p = 0.018; t _cluster_ (24) = 19.52, p = 0.002) during MW relative to BF (see Figure 4B). Again, a qualitatively similar pattern of results was obtained with a stricter definition of peaks (using the 1/trend of the spectrum as a threshold) (see Supplementary Figure 3B) (t _cluster_ (24) = 30.27, p = 0.003; t _cluster_ (24) = 19.12, p = 0.016).

### Condition-related modulations of alpha:theta phase synchrony

Permutation statistics revealed a significant increase in alpha:theta phase synchrony (as quantified by 2:1 and 3:1 phase locking value (PLV)) during MW relative to BF condition. One significant positive cluster was identified for 2:1 phase synchrony (t _cluster_ (24) = 8.43, p = 0.016) (Figure 5A) whilst two significant positive clusters were identified for 3:1 phase synchrony (t _cluster_ (24) = 27.79; p= 0.020; t _cluster_ (24) = 7.43; p = 0.021) (Figure 5B). A qualitatively similar pattern of results was obtained when estimating phase synchrony only in those trials showing alpha and theta peaks above an estimation of the 1/f trend of the power spectrum (2:1 PLV: t _cluster_ (24) = 10.17, p = 0.073) (3:1 PLV: t _cluster_ (24) = 3.03, p = 0.004; t _cluster_ (24) = 6.51, p = 0.012) (see Supplementary Figure 4).

**Figure 5.**
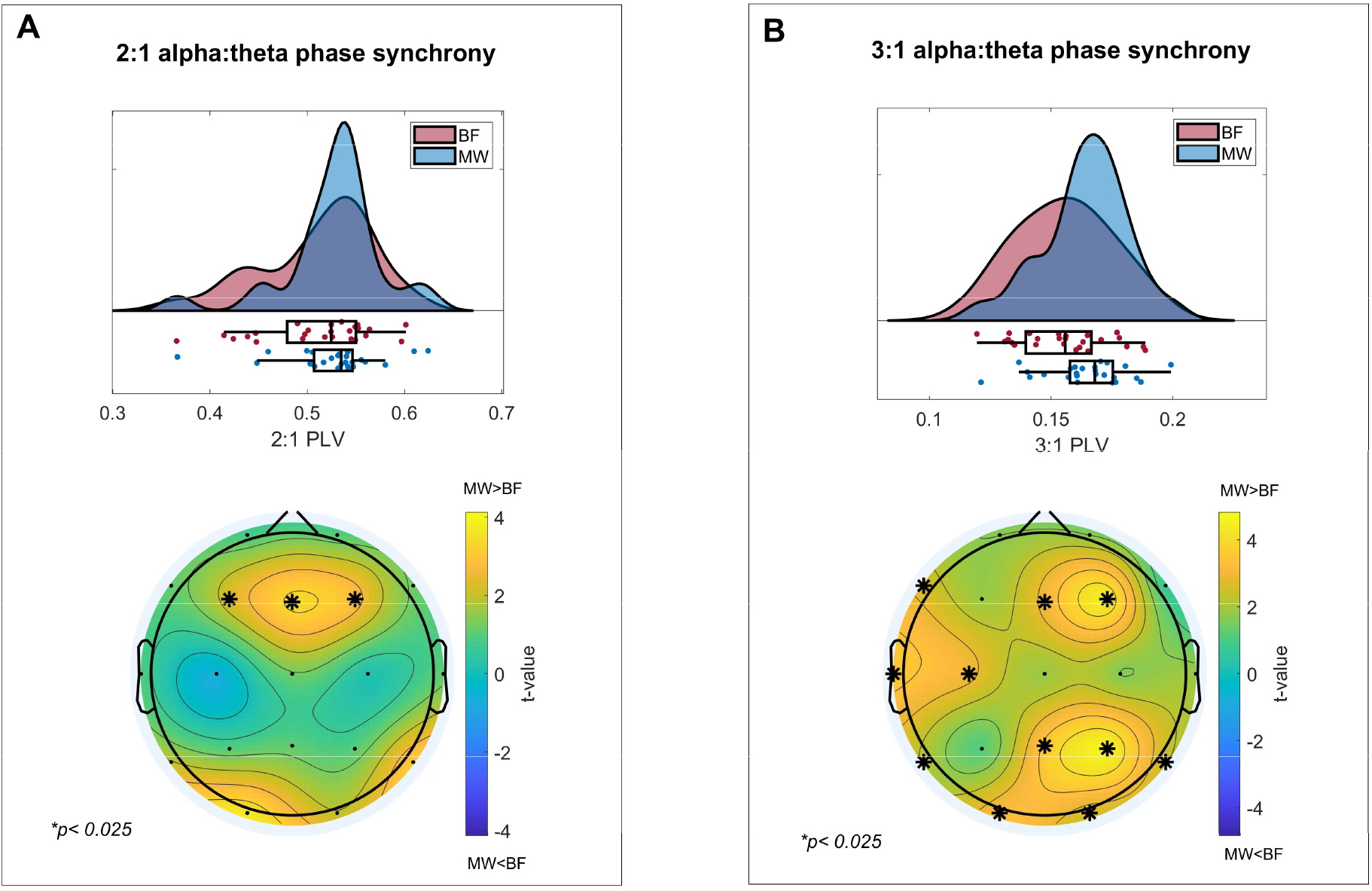
Condition-related modulations of alpha:theta cross-frequency phase synchrony. Each panel depicts the topographical maps representing condition-related changes (MW - BF) in each electrode (bottom plot; t-values of paired samples t-test are color-coded; asterisks mark cluster corrected p-values < 0.025) and the probability distributions of phase locking values (PLV) for the identified significant clusters in each condition (top plot; MW = mind wandering; BF = breath focus; individual points are subjects). Both 2:1 (Panel A) and 3:1 (Panel B) phase synchrony between alpha (8-14 Hz) and theta (4-8 Hz) rhythms were significantly increased during MW relative to BF.

### Debriefings of drowsiness, arousal and emotional valence

Retrospective debriefings (at the end of the task) showed that, on average, participants reported to feel drowsy approximately 50 % of the time during the task (mean Likert score = 3.82; SD = 1.52). Notably, the level of drowsiness was significantly correlated to the percentage of MW trials (r (20) = 0.61, p = 0.0022) indicating that participants with a higher proportion of probe-caught MW trials reported to feel more drowsy during the task. No significant correlations were found however, between the percentage of MW trials and inter-individual differences in retrospective debriefings of the participants’ level of arousal (r (20) = 0.09; p = 0.68) or emotional valence (r (20) = 0.03; p = 0.86). In addition, the level of arousal - as assessed with trial-specific debriefings - was not significantly different between MW and BF conditions (t (24) = −0.22, p = 0.82).

### Assessing the effect of drowsiness in condition-related modulations

Given the relative high scores of drowsiness in our sample and their correlation with the incidence of self-reported mind wandering, we explored whether drowsiness levels impacted the reported EEG condition-related differences. First, we reduced the dimensionality of EEG dependent variables that showed significant condition-related differences (i.e. MW versus BF) by averaging within the previously identified significant positive or negative clusters. Then, we performed one-way repeated measures ANCOVAs for each dependent variable with condition as within-subject factor (MW vs BF) and drowsiness (scale from 1 to 7; zero-centered) as a covariate. These additional analyses revealed that the condition effect in all of the studied EEG variables remained significant after controlling for drowsiness (see Supplementary Table 1). In addition, the main effect of drowsiness (i.e. correlation between drowsiness and dependent variable across conditions) was not significant for any of the here studied EEG parameters. Note however that some of the EEG dependent variables showed a ‘condition by drowsiness’ interaction, either significantly (i.e. relative amplitude theta: F (1,20)=11.29, p = 0.003; relative amplitude alpha: F (1,20) = 7.44 p = 0.013) or at trend-level (i.e. cross-frequency ratios and phase synchrony; F (1,20) > 3; p < 0.1). In this way, post-hoc Pearson correlations revealed that the condition effect (difference between MW and BF condition) in these EEG parameters was more pronounced for subjects that reported to be drowsier. For a comprehensive view of this secondary analysis (i.e. F-values, r-values, p-values and correlation plots) see Supplementary Table 1 and Supplementary Figure 5.

## DISCUSSION

In this study, we used an experience sampling approach to assess EEG spectral modulations in the theta-alpha range during mind wandering in the context of breath focus meditation in a sample of 25 novice meditation practitioners. Self-reported mind wandering episodes were associated with an increase in the amplitude of theta oscillations (~4-8Hz) and a decrease in the amplitude of alpha oscillations (~8-14 Hz) relative to breath focus states. In addition, these amplitude modulations during self-reported mind wandering were accompanied by changes in mean frequency (i.e. higher mean alpha frequency and lower mean theta frequency) and cross-frequency dynamics between alpha and theta rhythms (i.e. increased 2:1/3:1 harmonicity and cross-frequency phase synchrony).

Alpha desynchronization and theta synchronization have been related to a wide variety of cognitive tasks involving retrieval, storage and/or manipulation of information (Klimesch, 1999; Klimesch et al., 2008; Sauseng et al., 2009). Previous literature suggests that alpha is responsible for the representational component (i.e. storage/accessibility of task relevant information) whilst theta subserves the executive one (i.e. retrieval/manipulation of information) (Akiyama et al., 2017; de Vries et al., 2020; Klimesch, 2012; Sauseng et al., 2010; Schack et al., 2005). It can be speculated that alpha desynchronization/theta synchronization during mind wandering reflects the retrieval and manipulation of memory templates (de Vries et al., 2020). This is in line with previous literature pointing to a key role of areas involved in memory and executive control in mind wandering (Ellamil et al., 2016; Fox et al., 2015). Hence, our study supports the notion that mind wandering could be integrated into executive models of attention (see Smallwood & Schooler, 2006, 2015) since it shares some neural correlates with controlled cognitive processing (Baddeley, 1996, 2010).

Two previous studies reported alpha desynchronization/theta synchronization during mind wandering when compared to breath focus in novice meditation practitioners (Braboszcz & Delorme, 2011; van Son et al., 2019). Importantly, these studies assessed self-caught mind wandering (i.e. pressing a button whenever participants became aware of their mind wandering) instead of probe-caught mind wandering (i.e. probing the participants at random time intervals to report whether they were mind wandering). Thus, our results suggest that mind wandering during meditation with and without awareness could share some neural correlates in novice practitioners (i.e. alpha desynchronization/theta synchronization). In this line, it is also important to underline that one previous study that assessed probe-caught mind wandering during mantra meditation in experienced practitioners found a decrease in both alpha and theta amplitude during mind wandering (Brandmeyer & Delorme, 2018). We speculate that trials that were compared to mind wandering in Brandmeyer and Delorme (2016) corresponded to deeper meditative states that are qualitatively different from breath focus states in novice practitioners. In fact, deep meditative states in experienced practitioners have been linked to the emergence of high amplitude theta (4-8 Hz) oscillations (Fell et al., 2010; Hebert & Lehmann, 1977; Lomas et al., 2015). Therefore, and in line with Fell, Axmacher, & Haupt (2010), we hypothesize that different stages of meditation practice are related to different neural oscillatory correlates and that an increase in theta amplitude would only occur in advanced stages. In this way, future studies with experienced meditation practitioners are needed to determine whether theta oscillations emerging during deep meditative states and mind wandering differ in their spatiotemporal characteristics.

In addition to the analysis of amplitude in the EEG theta-alpha range (4-14Hz), we adopted a cross-frequency approach in which power modulations are analysed and interpreted as changes in the frequency architecture that influence the interaction between different brain rhythms (Rodriguez-Larios, Faber, et al., 2020; Rodriguez-Larios, Wong, et al., 2020; Rodriguez-Larios & Alaerts, 2019). This analytical approach is based on a recent theory positing that modulations in the center frequency of different brain rhythms determine cross-frequency communication (Klimesch, 2013, 2018). In particular, harmonic/non-harmonic cross-frequency arrangements are anticipated to enable/disable cross-frequency interactions by affecting their phase synchronization (and therefore, information exchange; see Fries, 2015; Palva & Palva, 2017; Varela et al., 2001). In line with this theory, both an increased occurrence of harmonic relationships and enhanced cross-frequency phase synchrony between alpha and theta rhythms have been observed during cognitive tasks in which these two rhythms are thought to interact (i.e. tasks requiring retrieval, storage and manipulation of information) (Kawasaki et al., 2010; Rodriguez-Larios & Alaerts, 2019; Schack et al., 2005). On the other hand, recent work from our group showed that alpha-theta harmonicity during meditation: (i) is reduced when compared to non-meditative rest in experienced practitioners (Rodriguez-Larios, Faber, et al., 2020); and (ii) is negatively correlated to meditation training in novices (Rodriguez-Larios, Wong, et al., 2020). Linking these findings to the observed increases in harmonicity during controlled cognitive processes (Rodriguez-Larios & Alaerts, 2019), it was hypothesized that the observed reductions in the occurrence of cross-frequency harmonicity during meditaton practice are reflective of a reduced engagement of cognitive processes involved in mind wandering. The current study supports this hypothesis by showing that mind wandering during meditation practice is related to increased 2:1/3:1 harmonicity and cross-frequency phase synchrony between alpha and theta rhythms. Hence, in light of the present pattern of results and previous literature, we speculate that when retrieval, storage and manipulation of information is required (during mind wandering or controlled cognitive processes) alpha and theta rhythms separate from each other in the frequency spectrum to enable transient 2:1/3:1 harmonic cross-frequency ratios and therefore, neural communication (via cross-frequency phase synchrony). On the contrary, states in which participants experience focused attention to body sensations and a reduced engagement in spontaneous self-generated thought (i.e. mind wandering) are anticipated to entail the convergence of alpha and theta rhythms in the upper theta/lower alpha band because this frequency arrangement would avoid their cross-frequency phase synchrony by fostering their synchronization at the same frequency (ratio ~ 1) and/or their desynchronization via non-harmonic irrational ratios (e.g. ratios ~ 1.6; see Pletzer, Kerschbaum, & Klimesch, 2010). In this way, it is important to underline that focused attention and/or mind wandering states during meditation practice might be experienced qualitatively differently in expert and novice practitioners. Although the current literature does not suggest that these differences are markedly reflected in alpha-theta cross-frequency dynamics (Rodriguez-Larios, Faber, et al., 2020; Rodriguez-Larios, Wong, et al., 2020), future studies are needed to directly assess this possibility. In this way, an EEG study adopting an experience sampling paradigm with both novices and experienced practitioners could clarify whether the here reported modulations in alpha—theta cross-frequency dynamics during mind wandering versus focused attention states differ between these two groups.

An important limitation of our study is the possible influence of drowsiness in phenomena categorized as mind wandering by the participants. The majority of the participants reported a high degree of drowsiness during the breath focus task (i.e. ~ 50% of the time). Moreover, the condition effect in some of our EEG dependent variables (i.e. relative amplitude) seemed to significantly moderated by the level of drowsiness (i.e. more drowsy subjects showed a more pronounced condition effect). Although no significant moderation could be found for alpha-theta cross-frequency parameters, some statistical trends could be observed. Thus, we speculate that the here reported changes in alpha-theta (cross-frequency) dynamics could be related to hypnagogic states (i.e. dream-like states in the transition to sleep; Schacter, 1976) that could have been categorized as mind wandering by the participants. In order to control for the effect of drowsiness in those episodes categorized as mind wandering, future studies could include a drowsiness scale in each trial and exclude those trials with high levels of drowsiness. Another possible solution could be to use the same paradigm with experienced meditators. Since meditation training has been shown to negatively correlate with drowsiness during meditation practice (Cahn et al., 2010), it can be anticipated that experienced meditators would feel less drowsy during the breath focus task employed in this study. In this way, it is important to take into account that hypnagogic states and mind wandering might be difficult to distinguish from a phenomenological perspective (even for experienced practitioners). In fact, several authors have proposed that the neurophenomenology of mind wandering and dream-like states might differ quantitatively instead of qualitatively (Domhoff & Fox, 2015; Fox et al., 2013).

Our results support growing evidence pointing to the key role of EEG alpha (8-14Hz) and theta (4-8Hz) rhythms during mind wandering in the context of meditation practice (Braboszcz & Delorme, 2011; Brandmeyer & Delorme, 2018; van Son et al., 2019). Therefore, modulations in alpha and theta rhythms could serve as an adequate parameter for the development of EEG neurofeedback protocols aimed to facilitate meditation practice by the real-time detection of mind wandering episodes (Brandmeyer & Delorme, 2013). In this way, further research is needed to determine which alpha-theta parameter (amplitude, frequency, cross-frequency interaction) or combination of parameters is most suited to use for EEG-neurofeedback when taking factors like signal to noise ratio, trainability and speed of online processing into account. In this way, we believe that this future research is of crucial importance since EEG-guided meditation could be highly beneficial for clinical (e.g. autism spectrum disorder, attention deficit disorder, mood disorders…etc.) and non-clinical populations that may show special difficulties to notice and disengage from mind wandering episodes when necessary (Biederman et al., 2019; Killingsworth & Gilbert, 2010).

In conclusion, our study shows that self-reported mind wandering during breath focus meditation in novice practitioners is associated with EEG spectral changes in the theta-alpha frequency range. In particular, mind wandering (relative to breath focus) was associated with: (i) increased amplitude and decreased frequency of theta (4-8 Hz) oscillations, (ii) decreased amplitude and increased frequency of alpha oscillations (8-14 Hz), and (iii) increased harmonicity and phase synchrony between alpha and theta rhythms. Because similar EEG spectral changes have been reported during controlled cognitive processes involving memory and executive control (de Vries et al., 2020; Haegens et al., 2014; Kawasaki et al., 2010; Rodriguez-Larios & Alaerts, 2019; Sauseng et al., 2005), we speculate that higher order neurocognitive mechanisms involved in both self-generated thought and controlled cognitive processing could be minimized during states of focused attention to body sensations. From a translational perspective, this study suggests that theta and alpha rhythms could be key to develop EEG-neurofeedback protocols aimed at facilitating the practice of meditation by helping practitioners to gain awareness of the emergence of self-generated thoughts.

## ACKNOWLEDGEMENTS

We would like to thank Eduardo Bracho Montes de Oca, Savina de Radiguès and Marie Drion for their assistance during data collection.

This work was supported by the Branco Weiss fellowship of the Society in Science–ETH Zurich, Grants from the Flanders Fund for Scientific Research (FWO projects KAN 1506716N and G079017N) and the European Varela Awards (Mind & Life Europe).

## AUTHOR CONTRIBUTIONS

J.R. formulated the hypothesis, organized the data collection, performed the analysis and drafted the first version of the manuscript. J.R. and K.A. conceptualized the study and reviewed / approved the final manuscript.

## DATA ACCESIBILITY

Raw data and MATLAB code will be publically available after publication in the Open Science Framework webpage (see https://osf.io/b6rn9/?view_only=b260d7611a664e87a3c291bcd9367a7b).

## SUPPLEMENTARY MATERIALS

**Supplementary Figure 1.**
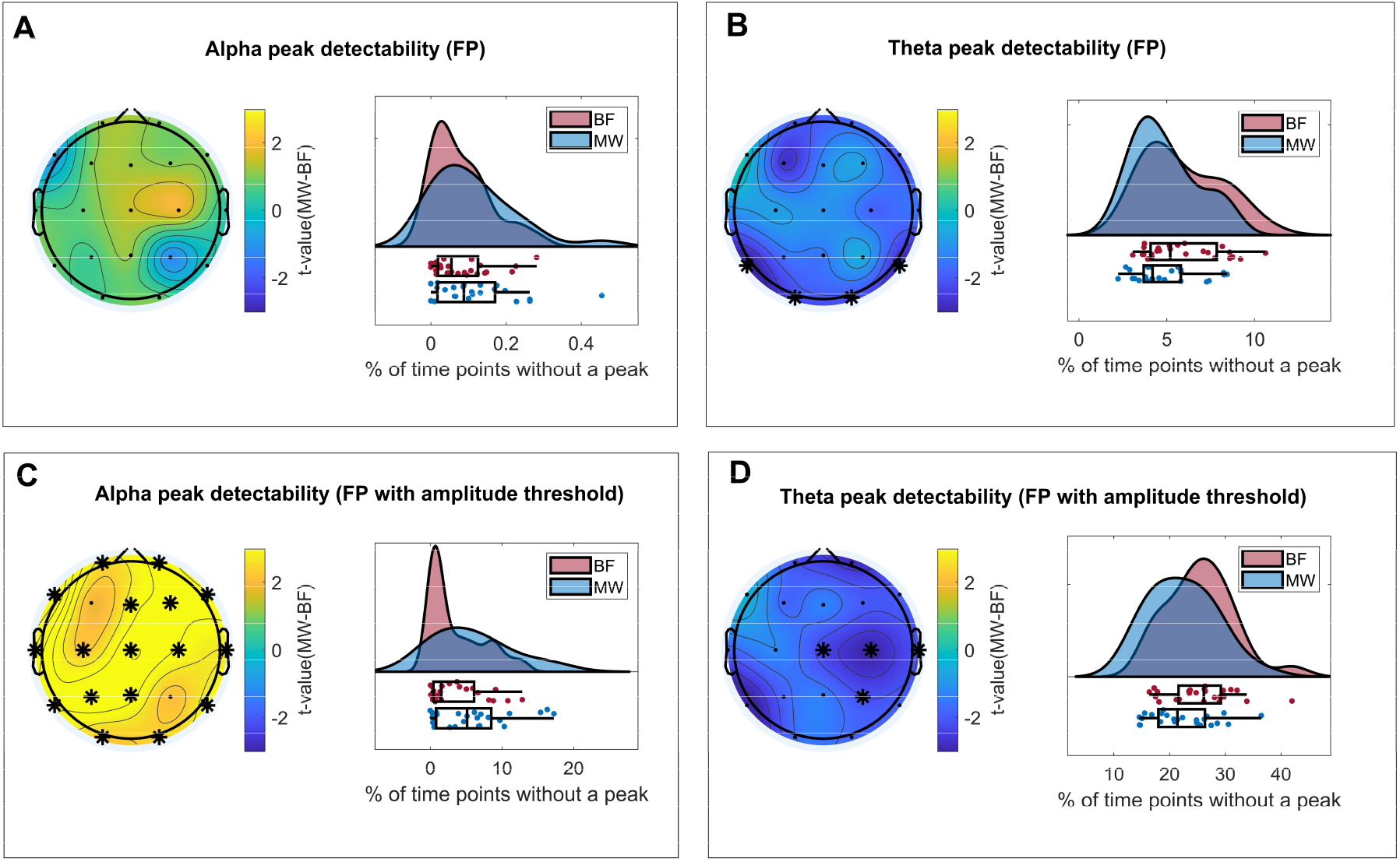
Condition-related modulations of peak detectability in alpha and theta frequency bands. Each panel depicts topographical maps representing condition-related changes in mean alpha (8-14 Hz) and theta (4-8 Hz) peak detectability (mind wandering (MW) - breath focus (BF)) for each electrode (left plot; t-values of paired samples t-test are color-coded) and their probability distributions per condition (right plot; % of time points without a peak averaged across electrodes). When estimated without amplitude threshold (Panel A and B), MW was associated to higher theta peak detectability (lower % of time points without a peak relative to BF). When peak detectability was computed with an amplitude threshold (estimation of 1/f trend; Panel C and D), MW was associated with higher theta peak detectability and lower alpha peak detectability (relative to BF).

**Supplementary Figure 2.**
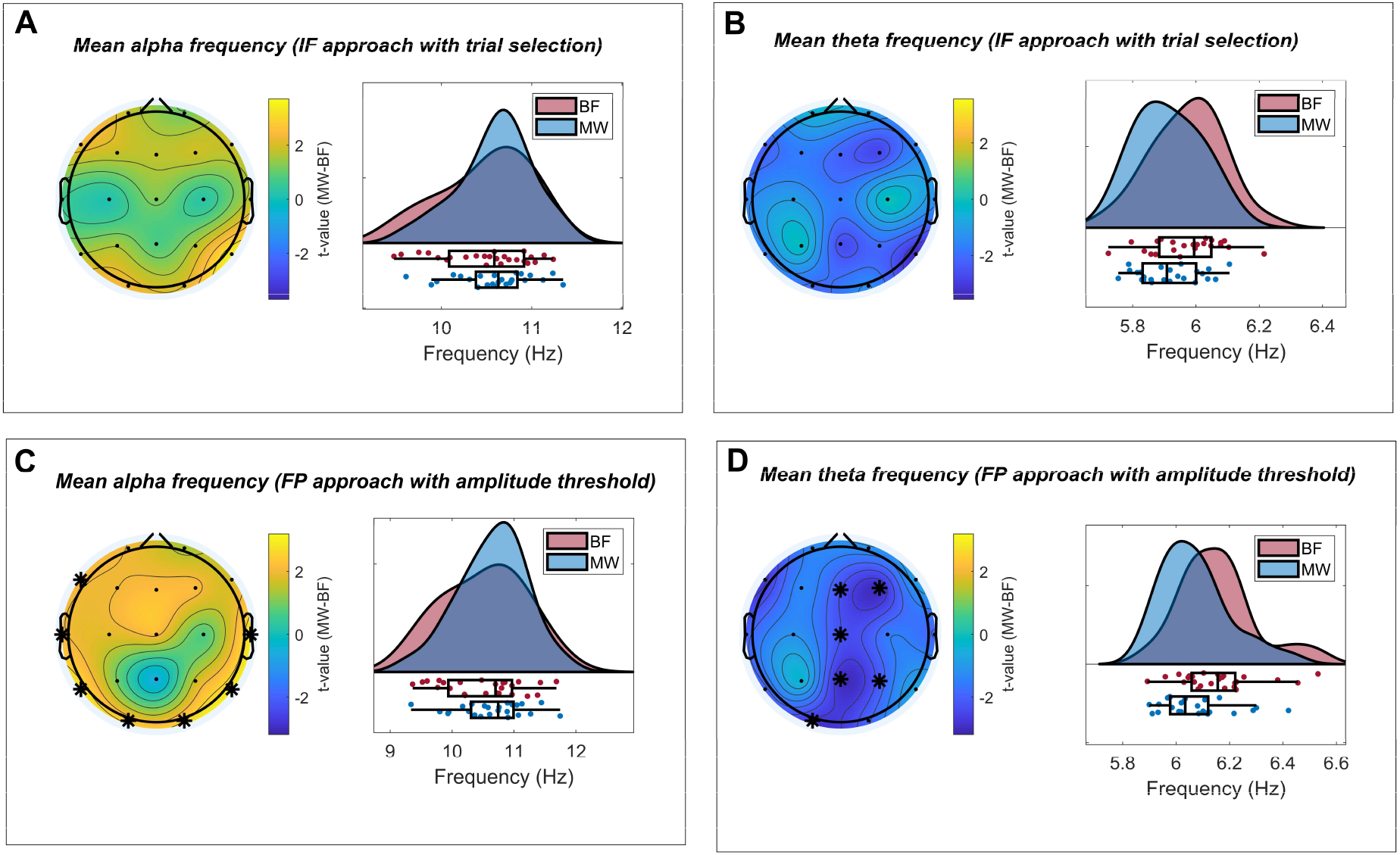
Condition-related modulations of alpha and theta mean frequency. Each panel depicts topographical maps representing condition-related changes (mind wandering (MW) - breath focus (BF)) in mean alpha and theta frequencies per electrode (left plot; t-values of paired samples t-test are color-coded) and the distributions of the identified clusters per condition (right plot; individual points are subjects). Mean frequency in alpha (8-14 Hz) and theta (4-8 Hz) bands were estimated through instantaneous frequency (IF) (selection of trials with alpha and theta peaks above the 1/f trend of the spectrum) and find peaks (FP) (amplitude threshold based on the 1/f trend of the spectrum) analytical approaches. When estimated with the instantaneous frequency (IF) approach, MW (relative to BF) was associated with a trend-level increase in mean alpha frequency (Panel A) and a trend level decrease in mean theta frequency (Panel B). A qualitatively similar pattern of results could be observed when mean frequencies were estimated with the FP approach (Panel C and D), although in this case, these changes did reach statistical significance (p cluster < 0.025).

**Supplementary Figure 3.**
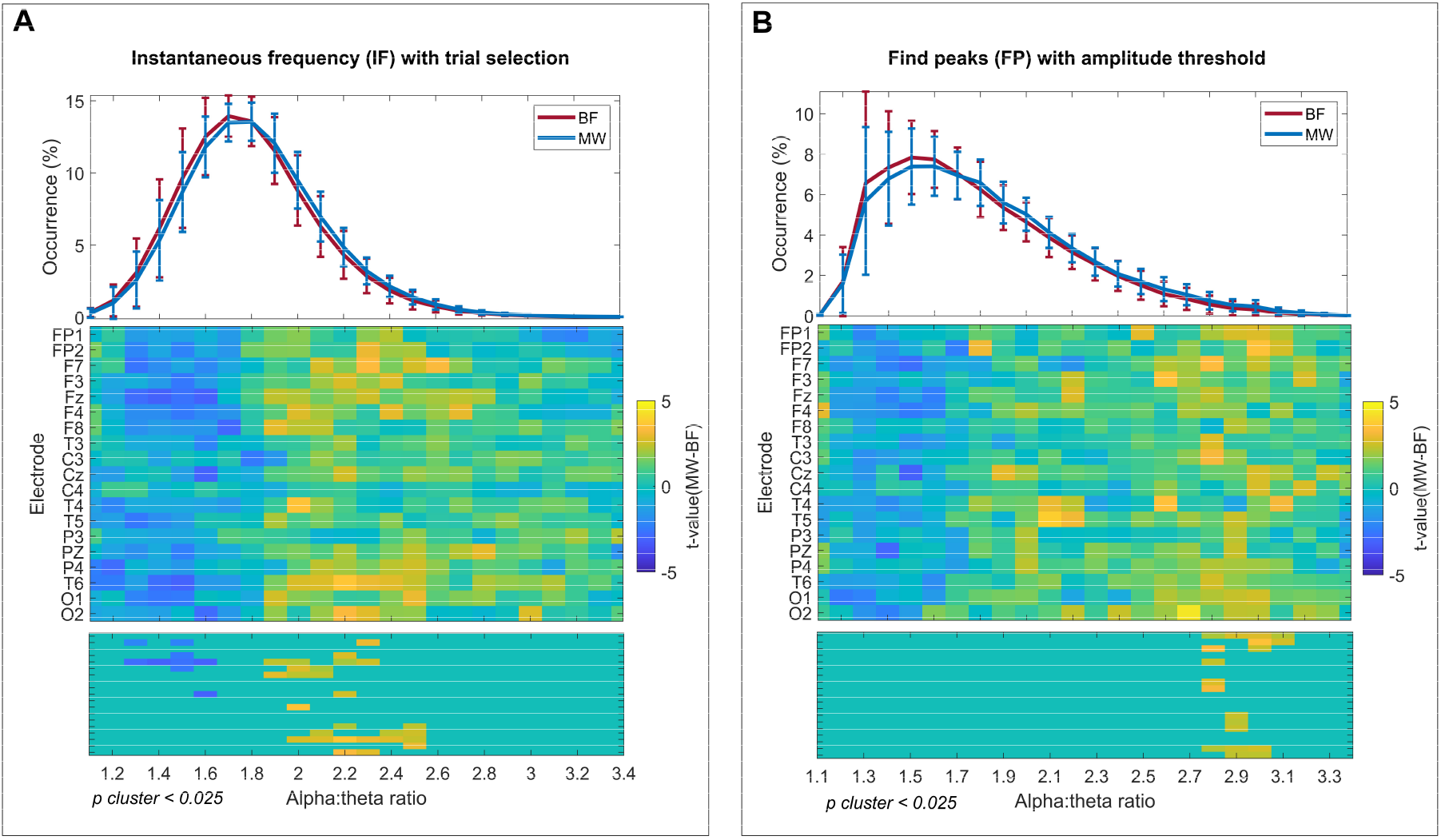
Condition-related modulations of transient alpha:theta cross-frequency ratios. Depiction of grand averages and condition effects (mind wandering (MW) – breath focus (BF)) in the occurrence of alpha:theta cross-frequency ratios when estimated with the instantaneous frequency (IF) (selection of trials with alpha and theta peaks above the 1/f trend of the spectrum) (Panel A) and find peaks (FP) (amplitude threshold based on the 1/f trend of the spectrum) (Panel B) analytical approaches. The top panels show the mean occurrence of each alpha:theta ratio (from 1.1 to 3.4 in steps of 0.1 Hz) per condition (MW in blue and BF in red) across electrodes and subjects (error bars represent standard deviation). Middle and bottom panels show t-values (resulting from paired samples t-tests) per cross-frequency ratio and electrode before and after correcting for multiple comparisons (cluster permutation correction in bottom plot; p-value of cluster < 0.025). When estimated through the IF approach, MW (relative to BF) was associated to an increase in the occurrence of alpha-theta cross-frequency ratios between 1.9 and 2.5 and a decrease in the occurrence of ratios between 1.3 and 1.6. For the FP approach, only a significant increase in the occurrence of alpha-theta ratios between 2.8 and 3.1 was reported.

**Supplementary Figure 4.**
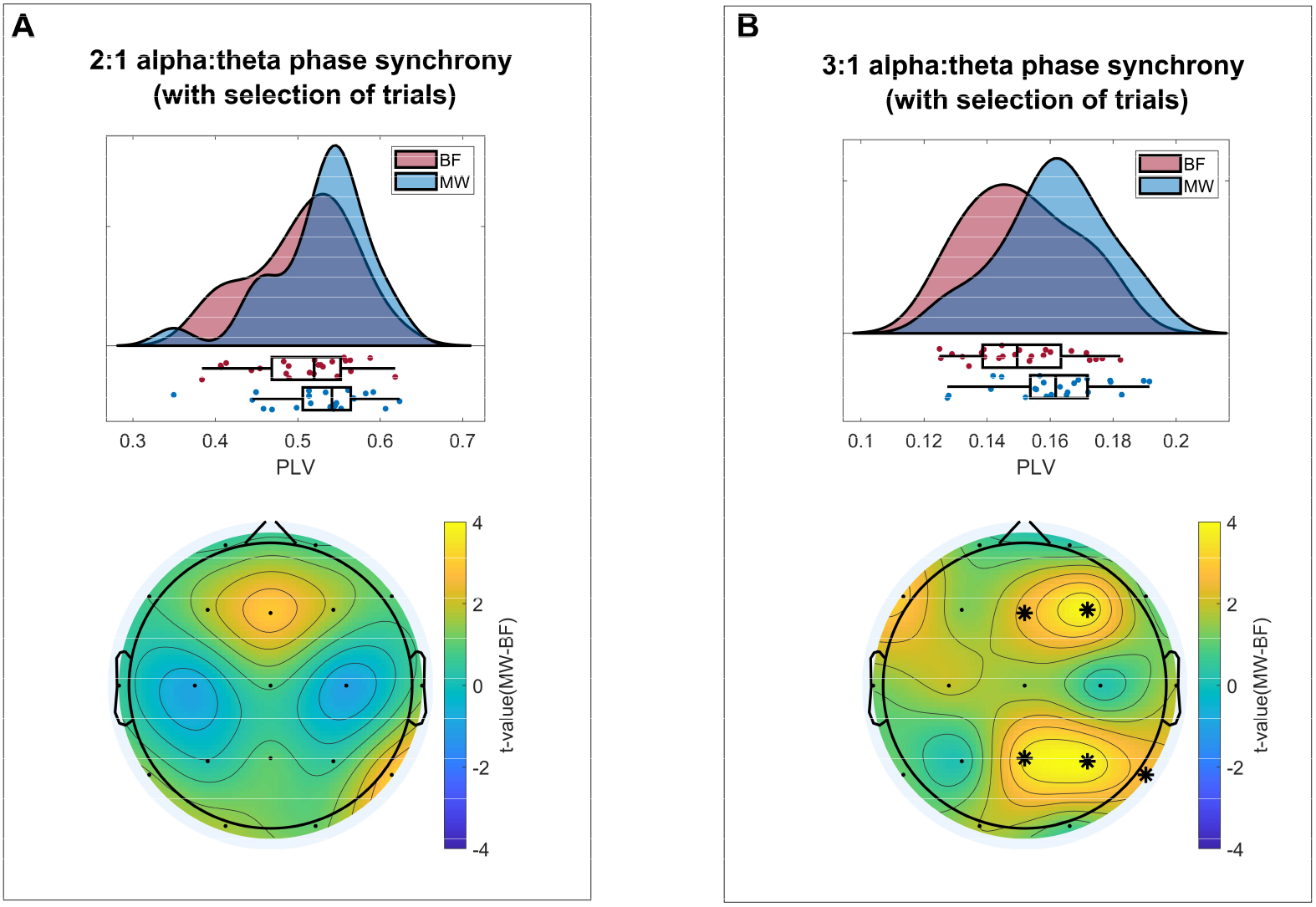
Condition-related modulations of alpha:theta cross-frequency phase synchrony. Each panel depicts the topographical maps representing condition-related changes (MW - BF) in each electrode (bottom plot; t-values of paired samples t-test are color-coded; asterisks mark cluster corrected p-values < 0.025) and the probability distributions of phase locking values (PLV) for the identified clusters in each condition (top plot; MW = mind wandering; BF = breath focus; individual points are subjects). For this analysis, only trials showing alpha and theta peaks above an estimation of the 1/f trend of the spectrum were taken into account. There was a trend level increase in 2:1 PLV (Panel A) and a significant increase in 3:1 PLV (Panel B) between alpha (8-14 Hz) and theta (4-8 Hz) rhythms during MW relative to BF.

**Supplementary Figure 5.**
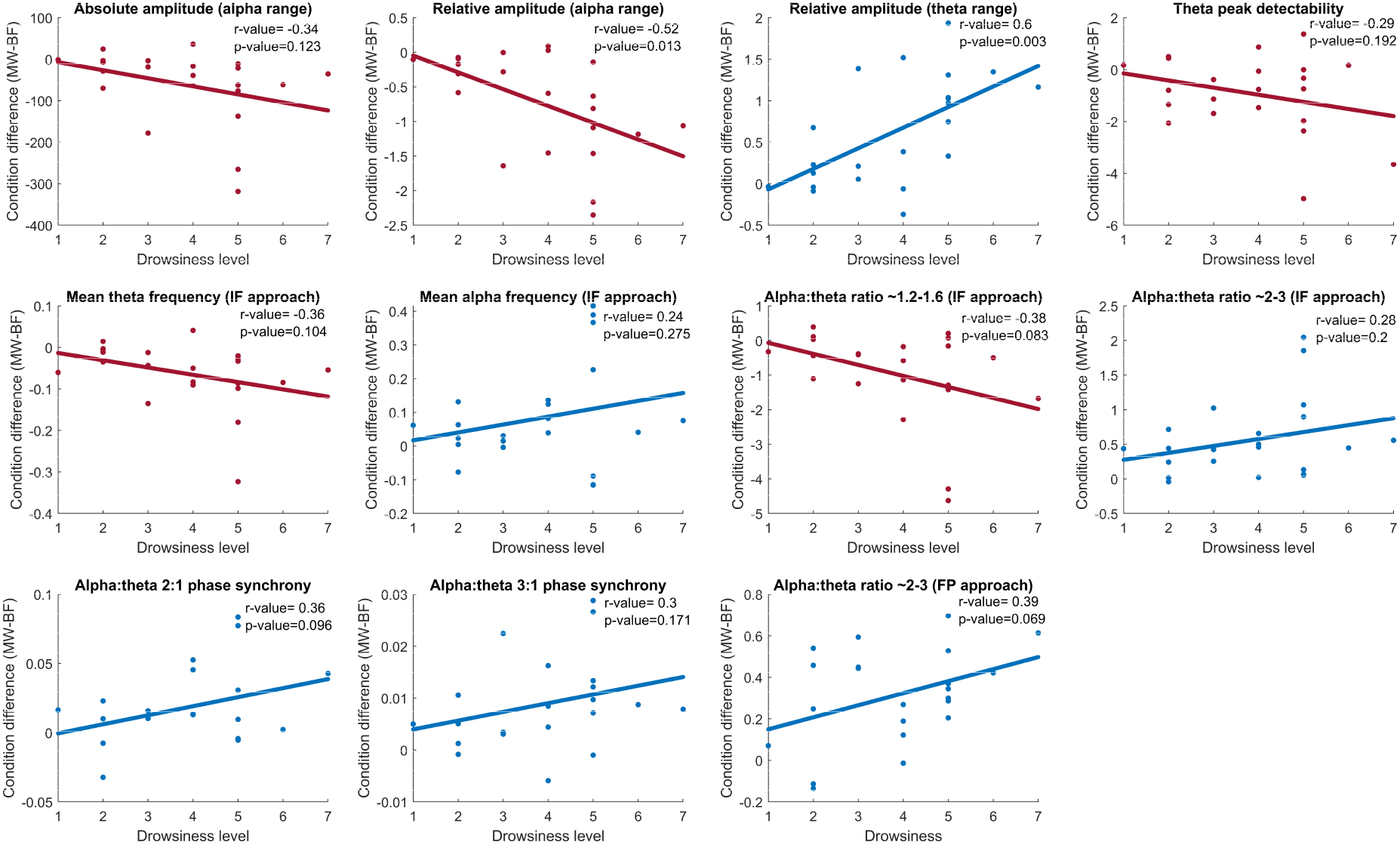
Correlations between condition-related modulations in different EEG parameters and drowsiness level. Each panel depicts the difference between mind wandering (MW) and breath focus (BF) conditions in a specific EEG parameter as a function of drowsiness level. Only EEG parameters that showed a significant condition effect were selected (averaged within positive or negative significant clusters). Individual points represent subjects. Pearson’s correlation coefficient and its corresponding p-value are shown in the top right corner of each panel. Significant correlations (indicating a more pronounced condition effect for subjects with higher drowsiness levels) were found for theta and alpha relative amplitude. Trend-level correlations were also identified for some cross-frequency parameters (i.e. 2:1 phase synchrony and cross-frequency ratios).

**Supplementary Table 1.**
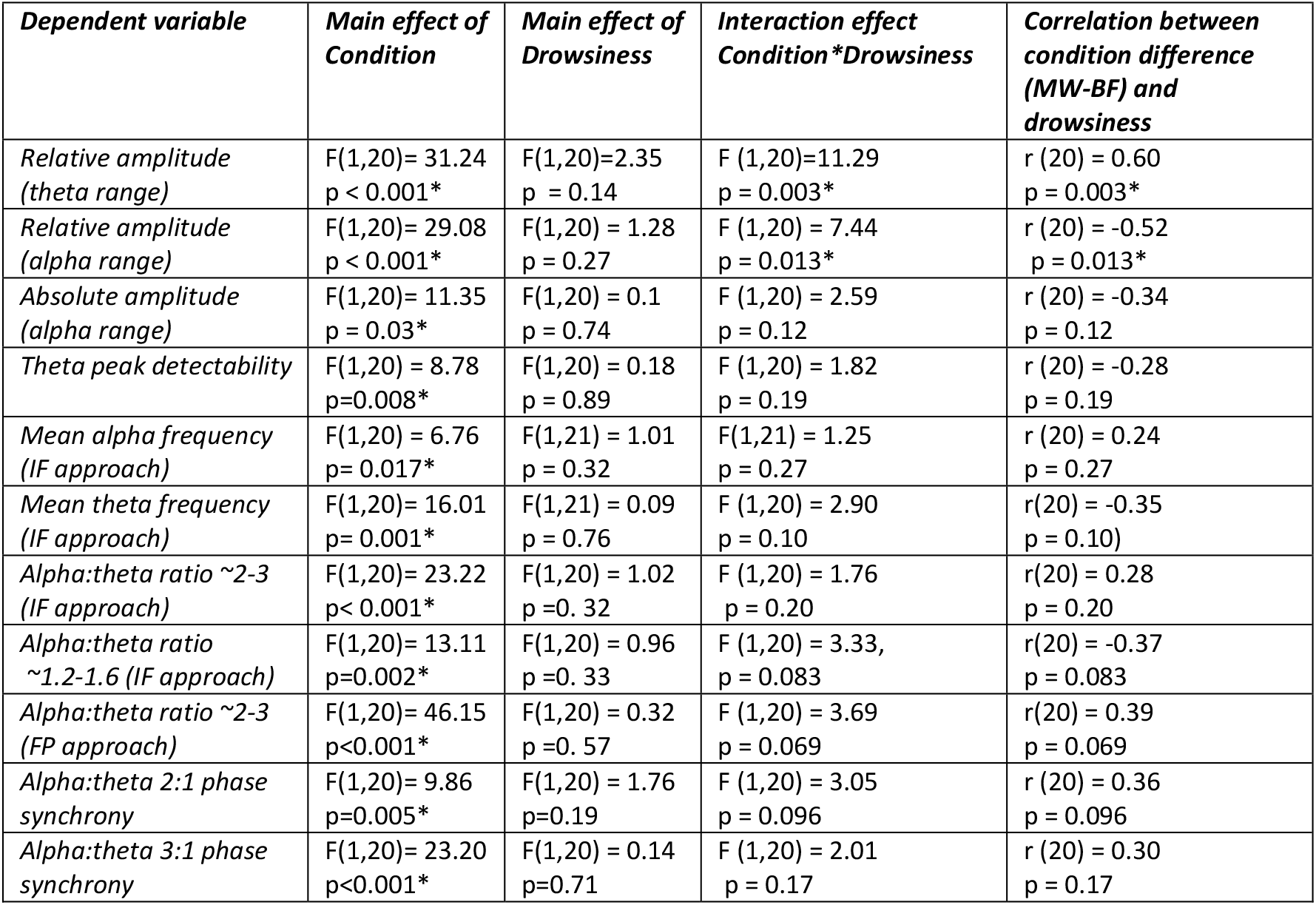
Results of repeated measures ANCOVAs assessing the effect of drowsiness in condition-related modulations. For the repeated measures ANCOVAs, condition (mind wandering (MW) and breath focus (BF) episodes) was set as a within-subject factor and drowsiness as a covariate. Post-hoc Pearson--correlation analyses are also reported, assessing the relationship between the condition-effect (difference between MW and BF conditions) and the level of drowsiness.

| *Dependent variable* | *Main effect of Condition* | *Main effect of Drowsiness* | *Interaction effect Condition\*Drowsiness* | *Correlation between condition difference (MW-BF) and drowsiness* |
| --- | --- | --- | --- | --- |
| *Relative amplitude (theta range)* | F(1,20)= 31.24<br>p < 0.001* | F(1,20)=2.35<br>p = 0.14 | F (1,20)=11.29<br>p = 0.003* | r (20) = 0.60<br>p = 0.003* |
| *Relative amplitude (alpha range)* | F(1,20)= 29.08<br>p < 0.001* | F(1,20) = 1.28<br>p = 0.27 | F (1,20) = 7.44<br>p = 0.013* | r (20) = -0.52<br>p = 0.013* |
| *Absolute amplitude (alpha range)* | F(1,20)= 11.35<br>p = 0.03* | F(1,20) = 0.1<br>p = 0.74 | F (1,20) = 2.59<br>p = 0.12 | r (20) = -0.34<br>p = 0.12 |
| *Theta peak detectability* | F(1,20) = 8.78<br>p=0.008* | F(1,20) = 0.18<br>p = 0.89 | F (1,20) = 1.82<br>p = 0.19 | r (20) = -0.28<br>p = 0.19 |
| *Mean alpha frequency (IF approach)* | F(1,20) = 6.76<br>p= 0.017* | F(1,21) = 1.01<br>p = 0.32 | F(1,21) = 1.25<br>p = 0.27 | r (20) = 0.24<br>p = 0.27 |
| *Mean theta frequency (IF approach)* | F(1,20)= 16.01<br>p= 0.001* | F(1,21) = 0.09<br>p = 0.76 | F (1,20) = 2.90<br>p = 0.10 | r(20) = -0.35<br>p = 0.10) |
| *Alpha:theta ratio ~2-3 (IF approach)* | F(1,20)= 23.22<br>p< 0.001* | F(1,20) = 1.02<br>p=0. 32 | F (1,20) = 1.76<br>p = 0.20 | r(20) = 0.28<br>p = 0.20 |
| *Alpha:theta ratio ~1.2-1.6 (IF approach)* | F(1,20)= 13.11<br>p=0.002* | F(1,20) = 0.96<br>p=0. 33 | F (1,20) = 3.33,<br>p = 0.083 | r(20) = -0.37<br>p = 0.083 |
| *Alpha:theta ratio ~2-3 (FP approach)* | F(1,20)= 46.15<br>p<0.001* | F(1,20) = 0.32<br>p=0. 57 | F (1,20) = 3.69<br>p = 0.069 | r(20) = 0.39<br>p = 0.069 |
| *Alpha:theta 2:1 phase synchrony* | F(1,20)= 9.86<br>p=0.005* | F(1,20) = 1.76<br>p=0.19 | F (1,20) = 3.05<br>p = 0.096 | r (20) = 0.36<br>p = 0.096 |
| *Alpha:theta 3:1 phase synchrony* | F(1,20)= 23.20<br>p<0.001* | F(1,20) = 0.14<br>p=0.71 | F (1,20) = 2.01<br>p = 0.17 | r (20) = 0.30<br>p = 0.17 |

